# State-dependent modulation of functional connectivity in early blind individuals

**DOI:** 10.1101/075465

**Authors:** Maxime Pelland, Pierre Orban, Christian Dansereau, Franco Lepore, Pierre Bellec, Olivier Collignon

## Abstract

Resting-state functional connectivity (RSFC) studies have highlighted how visual experience influences the brain’s functional architecture. Reduced RSFC coupling between occipital (visual) and temporal (auditory) regions has been reliably observed in early blind individuals (EB) at rest. In contrast, task-dependent activation studies have repeatedly demonstrated enhanced co-activation and connectivity of occipital and temporal regions during auditory processing in EB. To investigate this apparent discrepancy, the functional coupling between temporal and occipital networks at rest was directly compared to that of an auditory task in both EB and sighted controls (SC). Functional brain clusters shared across groups and cognitive states (rest and auditory task) were defined. In EBs, we observed higher occipito-temporal correlations in activity during the task than at rest. The reverse pattern was observed in SC. We also observed higher temporal variability of occipito-temporal RSFC in EB suggesting that occipital regions in this population may play a role of multiple demand system. Our study reveals how the connectivity profile of sighted and early blind people is differentially influenced by their cognitive state, bridging the gap between previous task-dependent and RSFC studies. Our results also highlight how inferring group-differences in functional brain architecture solely based on resting-state acquisition has to be considered with caution.

**Highlights:**

1. Occipito-temporal functional connectivity is modified by cognitive states.
2. This modulation is different in blind and sighted individuals.
3. Blind participants have higher occipito-temporal temporal variability at rest.
4. The group difference in variability at rest explains the differences in modulation.
5. Inferring group differences with resting-state data should be subject to caution.

## 1. Introduction

The study of people deprived of sensory information early in life provides conclusive evidence on how sensory experience shapes the structural and functional architecture of the brain (Frasnelli et al., 2011). Recent researches involving early blind individuals have shed new lights on the old ‘nature versus nurture’ debate regarding brain development: whereas the recruitment of occipital regions by non-visual inputs in the congenitally blind highlights its dependence upon experience to organize itself (nurture’s influence), the observation of specialized cognitive modules in the occipital cortex of congenitally blind, similar to those observed in the sighted, highlights the intrinsic constraints imposed to such plasticity (nature’s influence) (Collignon et al., 2011; Dormal and Collignon, 2011; Reich et al., 2012; Ricciardi et al, 2014).

Gaining deeper insights into how occipital regions in early blind individuals (EBs) are reorganized to support sensorimotor and cognitive functions not only requires studying their response properties (functional specialization), but also understanding how they are integrated into brain networks (functional integration) (Friston 2003). An increasingly popular approach to study short and long-range brain interactions is to measure functional connectivity (FC) during a resting-state (RS; van den Heuvel and Hulsoff Pol, 2010). With resting-state functional connectivity (RSFC), spontaneous fluctuations in functional magnetic resonance imaging (fMRI) signal, observed while participants are resting, are correlated to infer FC between intrinsically connected brain regions (Biswal et al., 1995). Therefore, RSFC is a measure of temporal synchrony of fMRI signal between distinct brain regions (Friston et al. 1993). Since functional networks have distinct temporal characteristics, separate functional networks can be identified from a single time series of resting fMRI data (Beckmann et al. 2005).

Research on the effect of early visual deprivations using RSFC protocols has evidenced both enhanced segregation (i.e. decreased connectivity) and integration (i.e. increased connectivity) of occipital regions (see Bock and Fine, 2014 for review). A striking and often replicated result emerging from those studies is the reduced connectivity of EBs’ occipital regions with primary somatosensory and auditory areas (Bedny et al., 2011; Burton et al., 2014; Liu et al., 2007; Yu et al., 2008; Striem-Amit et al., 2015; see Bock and Fine, 2014 for a recent review). This reliable effect at rest contrasts with the large amount of studies reporting enhanced task-dependent activations of occipital regions during the processing of auditory and tactile inputs (see Bavelier and Neville, 2002; Frasnelli et al., 2011 for review) in EBs. Importantly, the functional relevance of occipital activity to EB’s non-visual sensory/perceptual processing is supported by studies showing reduced performance to specific tactile and auditory tasks following the disruption of EB’s occipital cortex through the application of transcranial magnetic stimulation (TMS) (Cohen et al., 1997; Collignon et al., 2007, 2009). Moreover, TMS directly applied over the occipital cortex can elicit paresthesiae in the fingers of early blind braille readers (Cohen et al. 1997; Ptito et al., 2008). Further support comes from studies of effective connectivity (causal connectivity; Friston 1994) estimated by dynamic causal modelling of fMRI data (Friston 2003) or by combining TMS with Positron Emission Tomography (PET) (measuring activity induced by magnetically exciting distant cortical regions), which have shown stronger coupling between auditory or somatosensory areas and occipital regions in EBs than in SCs (fMRI: Collignon et al., 2013; Klinge et al., 2010, PET and TMS: Wittenberg et al., 2004). Finally, the participation of occipital cortex in non-visual processes early in time following stimuli presentation (Collignon et al., 2009; Leclerc et al., 2000; Röder et al., 1999) suggests the existence of direct links between occipital and other non-visual sensory regions.

We suggest that this apparent inconsistency between RSFC and task-dependent studies originates from the presupposition that RS networks are good proxies for networks instantiated during cognitive/perceptual processing (Andric and Hasson, 2015). Even though group level RSFC pattern have been documented to be similar to task co-activation patterns (Cordes et al., 2000; Smith et al., 2009; Wig et al., 2014), recent work has shown that whole-brain FC networks can be fundamentally reshaped by experience (e.g. Gordon et al., 2014; Orban et al., 2015; Lewis et al., 2009; Tambini et al., 2010), and over short periods of time (Allen et al., 2012; Hutchison et al., 2013b), that they are not constrained by RS topologies, and that this holds particularly true for connectivity structures of sensory systems (e.g. Andric and Hasson 2015; Mennes et al., 2013). Furthermore, investigations of the effect of task-focused attention versus rest/mind wandering in sighted individuals have shown functional coupling across brain regions to be state-dependent (Hampson et al., 2002; Hampson et al., 2004; Jiang et al., 2004; Newton et al., 2007). Specifically, functionally relevant connections show increased connectivity whereas irrelevant connections are decreased during task as compared to rest (Bartels and Zeki, 2005; Nir et al., 2006), with the resulting patterns of FC being specific enough to reliably predict participants' cognitive states (Shirer et al., 2012). Furthermore, evidence that differences in FC pattern between populations can be task-dependent (Çetin et al., 2014; Nair et al., 2014) compellingly illustrates how inferring differences in functional integration between groups solely based on RSFC should be subject to caution.

What is RSFC representative of then? The current view is that the wandering mind sequentially explores multiple functional modes, each with a unique pattern of connectivity, and sustaining a different function (Deco, Jirsa, & McInthosh, 2011; Karahanoğlu et al., 2013). Consequently, a connection’s strength will vary as a function of time (see Hutchison et al. 2013a,b), with more temporally variable connections being members of a larger set of modes (Allen et al., 2012). A secondary result of this dynamic nature is that connections measured over a whole run will be representative of no single functional mode, but of their average (Hutchison et al., 2013b, Smith et al., 2012). Of note, if a mode were to be less prevalent in a group, or in competition with a greater number of modes, its connectivity pattern would be more strongly diluted when computed over a whole run, and may consequently appear weaker.

Therefore, the goal of the current study was twofold. First, to investigate how cognitive states, rest and an auditory sensory/perceptual task, would impact FC between occipital and sensory cortices in both EBs and SCs. Second, to verify whether difference in RSFC between EBs and SCs is caused by their membership to a greater number of functional modes in the blind group. To investigate the topic of cognitive state, an auditory task was chosen because occipito-temporal RSFC has been reliably demonstrated to be lower in EBs than in SCs (Bedny et al., 2011; Burton et al., 2014; Liu et al., 2007; Striem-Amit et al., 2015; Yu et al., 2008), and because auditory processing elicits strong responses in both the occipital and temporal cortices of EBs (Collignon et al. 2011; 2013). Thus, participants underwent two fMRI runs, a RS one, and one where they were involved in a challenging auditory task. FC was extracted for each run and population, then the presence of an interaction between cognitive state and groups was tested. Additionally, for the RS run, a connection’s variability was measured as a function of time and the resulting metrics compared across groups. Biases caused by a priori definition of regions of interest (Zalesky et al. 2010; Park et al., 2013) were avoided by grouping voxels into functionally homogeneous regions. The actual method used, Bootstrap Analysis of Stable Clusters (Bellec et al., 2010), has recently been shown to perform well compare to other methods when it comes to subdividing the brain into meaningful functional regions (Ryali et al., 2015).

Based on studies revealing an increase in FC between functionally related areas and a decrease between functionally unrelated areas during tasks (Bartels and Zeki, 2005; Nir et al., 2006), our hypothesis was that the FC between auditory and occipital regions would be differently modulated by the cognitive states in EBs and SCs. Additionally, if some of EBs’ occipital regions do participate in a greater number of modes, their specific connections with auditory areas should be more variable than in SCs.

## 2 Materials and Methods

### 2.1 Participants

The data of fourteen EBs [4 females, age range 27–61 (mean ± SD, 42 ± 11] and 16 SCs [7 females, age range 23–60 (mean ± SD, 39 ± 14] were included in the analyses (see supplementary table 1 for more information on blind participants). Student’s t-test did not reveal any statistical age differences between groups. One additional SC participated in the study but was excluded from the analyses due to excessive motion during scanning acquisition (see below). Both groups were blindfolded throughout the fMRI acquisition. None of the EBs had ever had functional vision allowing pattern recognition or visually guided behaviour. At the moment of testing, all EBs were totally blind except for two who had only rudimentary sensitivity for brightness with no pattern vision. In all cases, blindness was attributed to peripheral deficits with no neurological impairment. For all subjects, pure-tone detection thresholds at octave frequencies ranging from 250 to 8,000 kHz were within normal limits in both ears. All of the procedures were approved by the research ethic and scientific boards of the Centre for Interdisciplinary Research in Rehabilitation of Greater Montreal and the Quebec Bio-Imaging Network. Experiments were undertaken with the understanding and written consent of each subject.

### 2.2 Experimental design

Participants underwent two functional runs. A RS acquisition was carried first in order to avoid task contamination of RS brain activity. This run lasted 5 minutes (136 volumes) during which participants were instructed to relax, not to think about anything in particular, and to keep their eyes closed.

The second run consisted of a 14 minutes long sound discrimination task (400 volumes; 278 acquired when participants are engaged the task) the method, behavioral results, and activation of which have already been described (Collignon et al. 2011; 2013). To summarize, the task run contained 30 blocks (20.4 s duration each) separated by rest periods. Participants had to process the spatial or pitch attributes of the sounds using a staircase method in order to keep correct responses at ~85% which allowed to equalize the difficulty level across tasks and participants. Since our goal was to assess the connectivity between auditory and visual regions independently of the specifics of the task, FC was equally based on data from the spatial and pitch blocks. The use of a staircase procedure inside the scanner notably allowed us to guarantee that the change in connectivity profile between groups is not due to the perceived difficulty level of the task.

### 2.3 FMRI data acquisition

Functional time series were acquired using a 3-T TRIO TIM system (Siemens) equipped with a 12-channel head coil. Multislice T2*-weighted fMRI images were obtained with a gradient echo-planar sequence using axial slice orientation [time to repetition (TR) 2,200 ms; time to echo (TE) 30 ms; functional anisotropy (FA) 90°; 35 transverse slices; 3.2-mm slice thickness; 0.8-mm interslices gap; field of view (FoV) 192 × 192 mm^2^; matrix size 64 × 64 × 35; voxel size 3 × 3 × 3.2 mm^3^]. A structural T1-weigthed 3D magnetization prepared rapid gradient echo sequence (voxel size 1 × 1 × 1.2 mm^3^; matrix size 240 × 256; TR 2,300 ms; TE 2.91 ms; TI 900 ms; FoV 256; 160 slices) was also acquired for all subjects.

### 2.4 FMRI data preprocessing

The datasets were analyzed using the NeuroImaging Analysis Kit (NIAK_0.7c3; http://niak.simexp-lab.org) (Bellec et al., 2012), under CentOS with Octave (v3.6.2; http://gnu.octave.org) and the Minc toolkit (v0.3.18-20130531; http://www.bic.mni.mcgill.ca/ServicesSoftware/ServicesSoftwareMincToolKit). All analyses were executed in parallel on the "Mammouth" supercomputer (http://www.rqchp.ca/fr/access-aux-ressources/serveurs/mp2), using the pipeline system for Octave and Matlab (PSOM 1.0) (Bellec et al., 2012).

Each fMRI run was corrected for inter-slice differences in acquisition time and rigid body motion. Then, the mean motion-corrected volume of functional data was coregistered with the T1 individual scan, which was itself non-linearly transformed to the Montreal Neurological Institute (MNI) non-linear symmetric template (Fonov et al., 2009; Fonov et al., 2011). Functional volumes were resampled to MNI space at a 3mm isotropic resolution. To minimize artifacts due to excessive motion, some time frames were removed (see method below). Afterward, slow time drifts (high-pass filter with a 0.01 Hz cut-off), the average signals in conservative masks of the white matter and the lateral ventricles, and the first principal components (95% energy) of six rigid-body motion parameters and their squares were regressed out of the time series. Finally, the volumes were spatially smoothed with a 6mm Gaussian kernel.

Absolute head motion parameters were compared between groups for both the task and RS datasets. No significant differences were found, neither between conditions and groups, nor for their interaction. To further rule out the possibility that the observed functional connectivity variability could be accounted for by inter-subject differences in head motion, the scrubbing method of Power et al. (2012) was used to remove volumes with excessive motion (frame displacement greater than 0.5 mm). A minimum number of 80 unscrubbed volumes per run, corresponding to 176s of acquisition, was then required for further analysis. For this reason, data from one SC were removed from the sample.

### 2.5 Functional brain clustering

The method developed by Bellec and colleagues (Bellec et al., 2010), called multi-level Bootstrap Analysis of Stable Clusters (BASC), was used to separate the brain into functional clusters for the connectivity analysis. Theses clusters each defined a circumscribed network or cerebral region. The choice for using BASC to obtain clusters was done with two goals in mind. First, it groups voxels into functionally homogeneous clusters across participants, a criteria usually not achieved through anatomical parcellation of the brain, and shown to affect FC results (Park et al., 2013). Second, the clustering solution of the method is representative of each group and condition since it is based on every functional time series that were recorded for this experiment Of note, consensus-based clustering such as BASC have been found to reliably identify functional subdivisions in a variety of brain areas with good biological plausibility and agreement with other imaging modalities (Ryali et al., 2015).

The first step of the analysis was to reduce the computational burden of the subsequent cluster analysis by deriving ~1,000 regions. To this end, a region-growing algorithm was applied on the voxelwise fMRI time series obtained from the concatenation of the rest and task runs (Bellec et al. 2006). These regions, called atoms, exclusively covered the grey matter and had a controlled size of 1000 mm^3^. Then, a cluster analysis was applied on the averaged fMRI time series of each atom to identify brain networks that consistently exhibited similar fluctuations between the rest and task runs in individual subjects.

At the group level, BASC quantifies the reproducibility of a cluster analysis, while providing a clustering solution that captures the most stable features across many bootstrap samples. To ascertain that the resulting clustering solutions would be equally representative of both groups (EBs; SCs) and cognitive states (rest; task), a stratified bootstrap sampling method, which creates samples that are balanced with respect to these factors, was implemented. Thus, functional clusters for this study were defined by consensus clusters, which were composed of regions with a high average probability of being assigned to a certain cluster in both groups and cognitive states. We generated a group-level consensus resolution of 50 clusters (see Fig. 1C) since this resolution provides an optimal trade-off between decreased sensitivity due to correction for multiple comparisons versus anatomical precision (Bellec et al., 2015). The center coordinates and average size of each cluster can be found in supplementary table 2 (see Fig. 2 for the clustering scheme).

**Figure 1:**
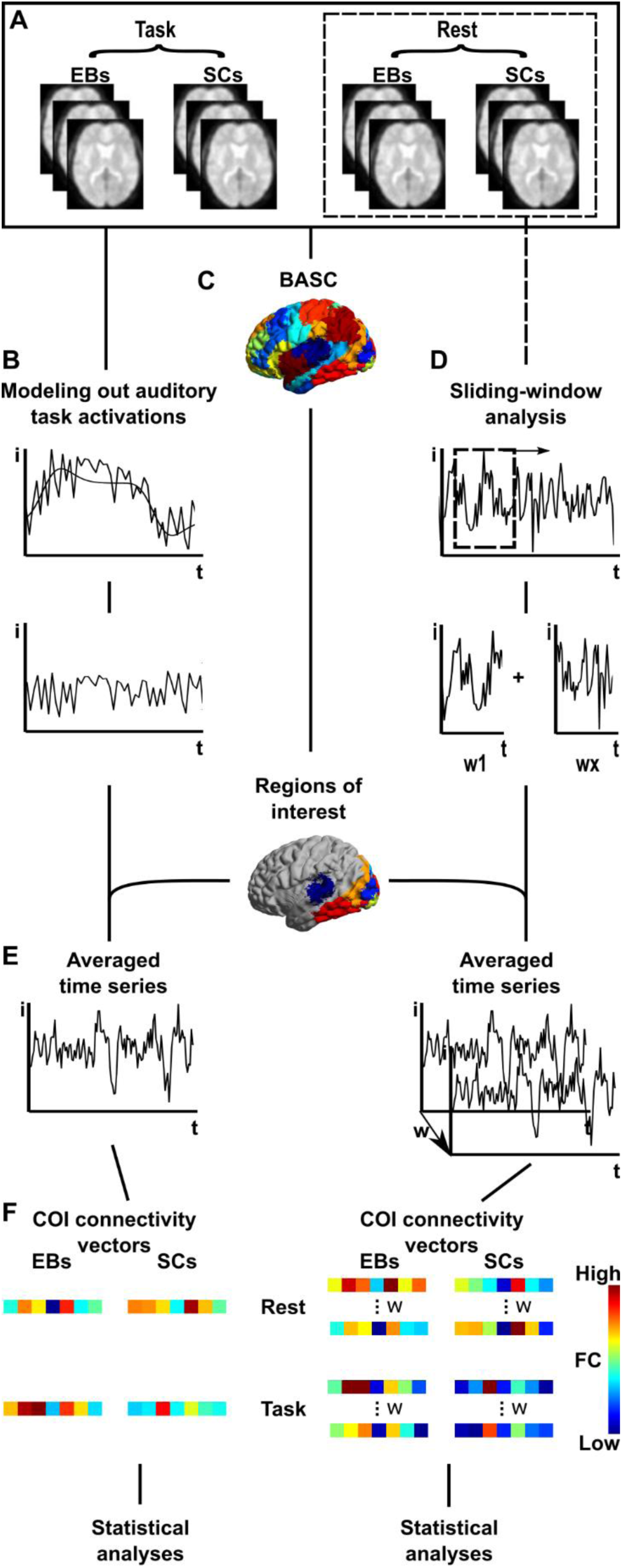
Steps of analyses. A) Preprocessed times series. B) Auditory task activations are modelled out of the task dataset. The modelisation is done at the voxel level independently for each participant and task. The resulting residuals are used as surrogate time series for the following analyses. C) Clustering of voxels using BASC. The clustering algorithm uses time series from both cognitive states and populations. D) Sliding-window analysis carried on RS data from EBs and SCs. Time series within windows of 21 TRs (broken line square) are extracted at the voxel level for each participant. This results in multiple time series per voxel per fMRI run (rest and task). E) Cluster averaged time series. Time series of all voxels within a cluster are. F) Computation of connectivity between the temporal cluster and the occipital ones. Graph axes are labeled with “i” for intensity, “t” for time, and “w” for time windows.

**Figure 2:**
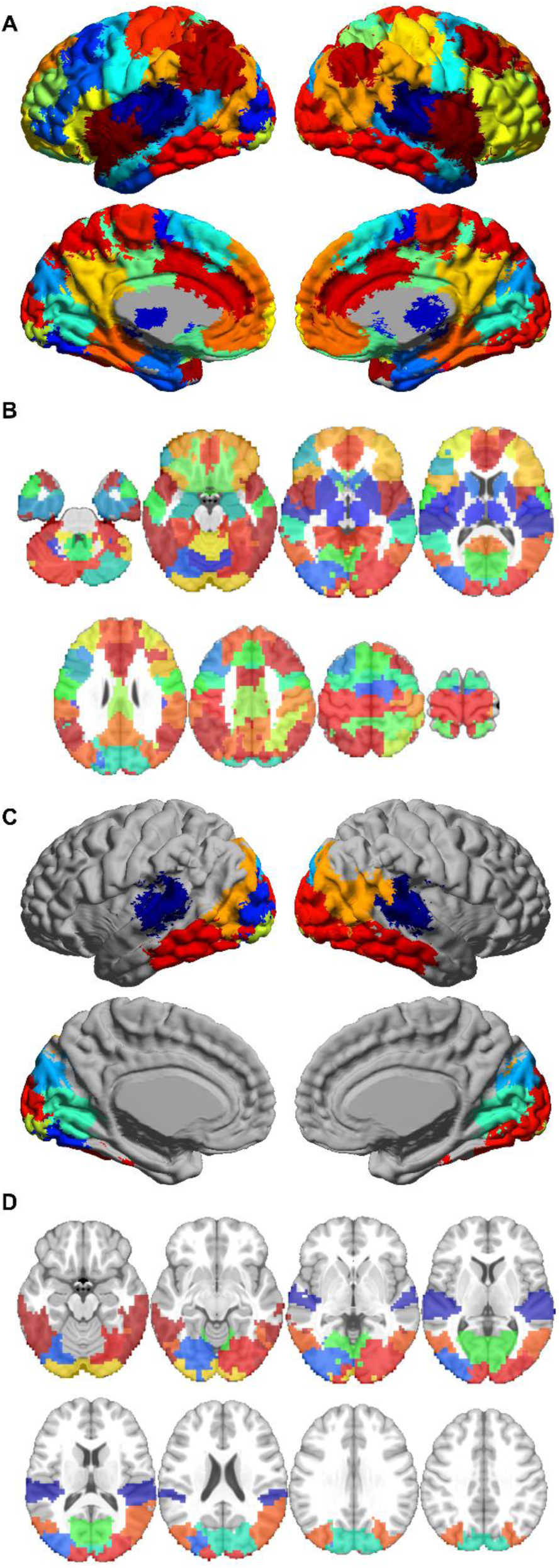
Results from the functional parcellation of the brain into 50 regions shown on a surface (A) and brain slices (B). Regions of interest selected for investigation of occipito-temporal FC, the temporal region and 7 occipital regions (D), are shown on a surface (C) and brain slices (D).

### 2.6 Occipito-temporal connectivity analysis

Based on our a priori hypothesis and the use of an auditory task, we focused our analyses to connections of interest (COI) between temporal (‘auditory’) and occipital (‘visual’) clusters. After separating the brain into 50 clusters, the auditory temporal cortex was limited to 1 bilateral cluster and the occipital cortex to 7 clusters (see Fig. 2C,D). Decisions as to which cluster should be considered part of the visual cortex was based on two criteria: 1) the most dorsal clusters were roughly limited by the parieto-occipital fissure, and 2) the most ventral clusters included the posterior fusiform gyri and the lingual gyri. Such liberal criteria were selected in order to include visual areas that are subject to neuroplastic changes following early blindness as indexed by unusual RS caracteristics (Liu et al., 2007; Liu et at., 2011; Burton et al., 2014) or functional reorganization following early visual deprivation (Collignon et al., 2011; Gougoux et al., 2005; 2009; Hölig et al., 2014).

Our connectivity analysis followed a 2 × 2 factorial design (groups × cognitive states), and was carried twice. In the first iteration, FC at rest was compared to FC during the task, the latter one being computed from time series which had had sustained task-evoked activity modeled out. In the second iteration, FC at rest was compared to FC during the task, but with the raw preprocessed time series. However, similarities between co-activations resulting from block designs and clusters linked by high FC (Cordes et al., 2000; Di et al., 2013; Laird et al., 2013; Smith et al., 2009; Wig et al., 2014) suggests that, for the task dataset, results obtained from raw preprocessed time series might be redundant with those provided by activation studies (Debas et al.2015). Thus, removing task-evoked sustained activity was necessary to avoid any FC effects which might arise from the blocked design of the auditory task (Fair et al. 2007; Nair et al., 2014) and provide results which are original. Regressing out task-evoked activity from time series was done by modeling task-related BOLD effects using a boxcar function that was convolved with the canonical hemodynamic function. Task-induced variance was removed from time series using the general linear model. The resulting residuals were used as surrogate time series for the computation of the auditory task FC.

Inter-cluster connectivity between every occipital clusters (n=7) and the temporal one (a single bilateral cluster) was defined as the Fisher transform of the Pearson correlation coefficient of the average time series of each cluster. The Fisher transform was implemented so that correlation coefficients would follow a Gaussian distribution (see Bellec et al., 2006). Of note, when cluster encompassed the two cerebral hemispheres, then its functional connectivity measure was based on the BOLD signal averaged across the two hemispheres. Since we were interested in comparing how FC is modified by the cognitive state in EBs and SCs, we then computed the modulation effect (which we define as the difference of FC between the two cognitive states: task FC–rest FC) for each connection and participant.

A random-effect group-level general linear model was estimated for the modulation effect of all COIs. Blindness, intercept, age, sex and motion, were entered as covariates with the later four being confounding variables. This allowed to test the presence of a main effect of groups (EB and SC) and to investigate whether some connections showed a stronger effect than others between groups (interaction of connections × groups). Effects of this omnibus test were considered significant for *p* < 0.05. Post-hoc tests were carried on single connections using the same general linear model as above. A false discovery rate (FDR) procedure (Benjamini and Hochberg 1995) was implemented to correct for the multiple comparisons arising from post-hoc tests. Threshold of statistical significativity was set at *q* < 0.05. Finally, since the intra-subject FC between any two connections was not significantly correlated in either the RS or task-state (Bonferonni or FDR corrected), no corrections were made for data dependency.

### 2.7 Functional connectivity variability analysis

A sliding window analysis was used to measure this variation over time. Since the removal of time frames by the scrubbing procedure would affect the temporal nature of the analysis, data were reprocessed using the same parameters as above but forgoing the scrubbing step. In order to still maximally control the effect of motion on FC, frame displacement (FD; Power et al., 2012) values were used as covariates during statistical analysis. Instead of using the full run, data from windows of 21 TRs (46.2 s) were employed. They were slid in steps of 1 TR resulting in a total of 116 windows per participant for the RS (similar to Allen et al., 2012, and considered to be optimal: Hendriks et al., 2016). Pearson's correlation coefficient between the time series of occipital and temporal clusters was computed for each window and then Fischer transformed. A functional connection’s variability across time was defined as the standard deviation of the aforementioned Fisher transformed correlation across time windows.

If some of EBs’ occipital regions do participate in a greater number of modes, their specific connections with auditory areas should be more variable than in SCs. This prediction was tested for connections of the COI analysis which had been found to have a different modulation effect in EBs than SCs. Each connection was individually tested using the GLM to control for variables of non-interest (age, sex, FD). Considering that only strong FC variability can be detected using fMRI runs as short as 5 minutes (Hindriks et al. 2015), any group differences unveiled by this analysis should be reliable.

To further investigate whether there exists a relationship between group differences in the modulation effects and group differences in a connection’s variability, we computed the average value of those variables for each occipito-temporal functional connections and correlated them. Our hypothesis was that connections showing a stronger modulation effect in EBs than SCs did so because they were involved in a larger number of networks and, hence, should be more temporally variable at rest in EBs.

## 3. Results

### 3.1 Occipito-temporal connectivity

As mentioned above, the COI analysis comparing the modulation effect across groups [EBs(task–rest) vs SCs(task–rest)] was run twice, once with surrogate time series from which the effect of the block design were removed, and once with raw preprocessed time series. Whether or not the effect of block was modeled out of the time series did not strongly affect FC values (see Supplementary table 3). Thus, only results for the tests carried on the time series from which task activations were modelled out are reported below.

An omnibus test [2 (groups) × 7 (connections) ANCOVA] showed no main effect of group (*p* < 0.11), but the interaction effect was significant (p < 0.001, Huynh-Feldt corrected) (Fig. 3), meaning that this effect was stronger for some connections than others. Post-hoc tests revealed two connections for which the between-group difference in modulation effect survived the FDR correction (Fig. 3 and 4). These connections involved the temporal region and the middle occipital gyrus (mostly the left hemisphere, #37) and the superior occipital gyrus (mostly the right hemisphere, # 47).

**Figure 3:**
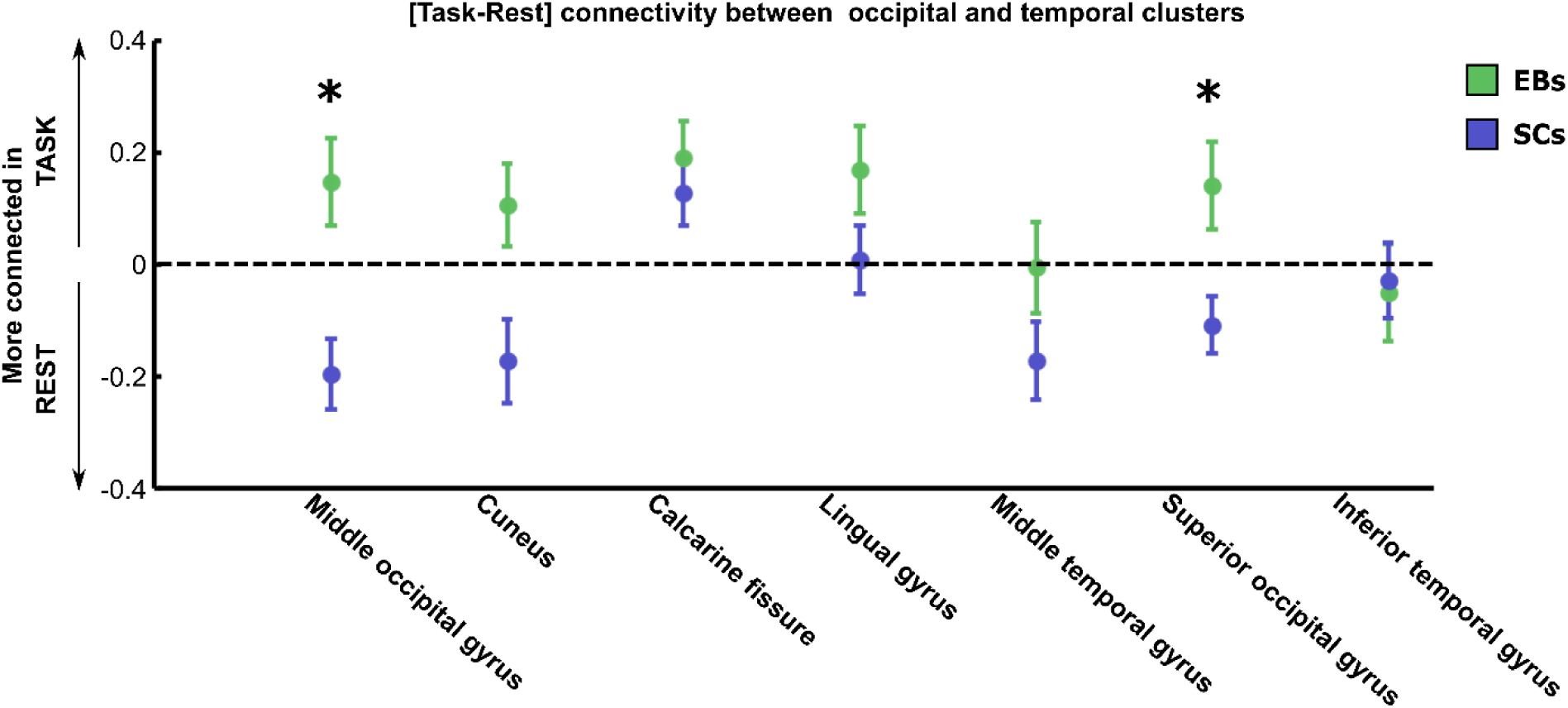
Group average and standard error of FC’s modulation effect [task – rest] for each connection between the temporal region and the 7 occipital regions. * q < 0.05 FDR corrected

**Figure 4:**
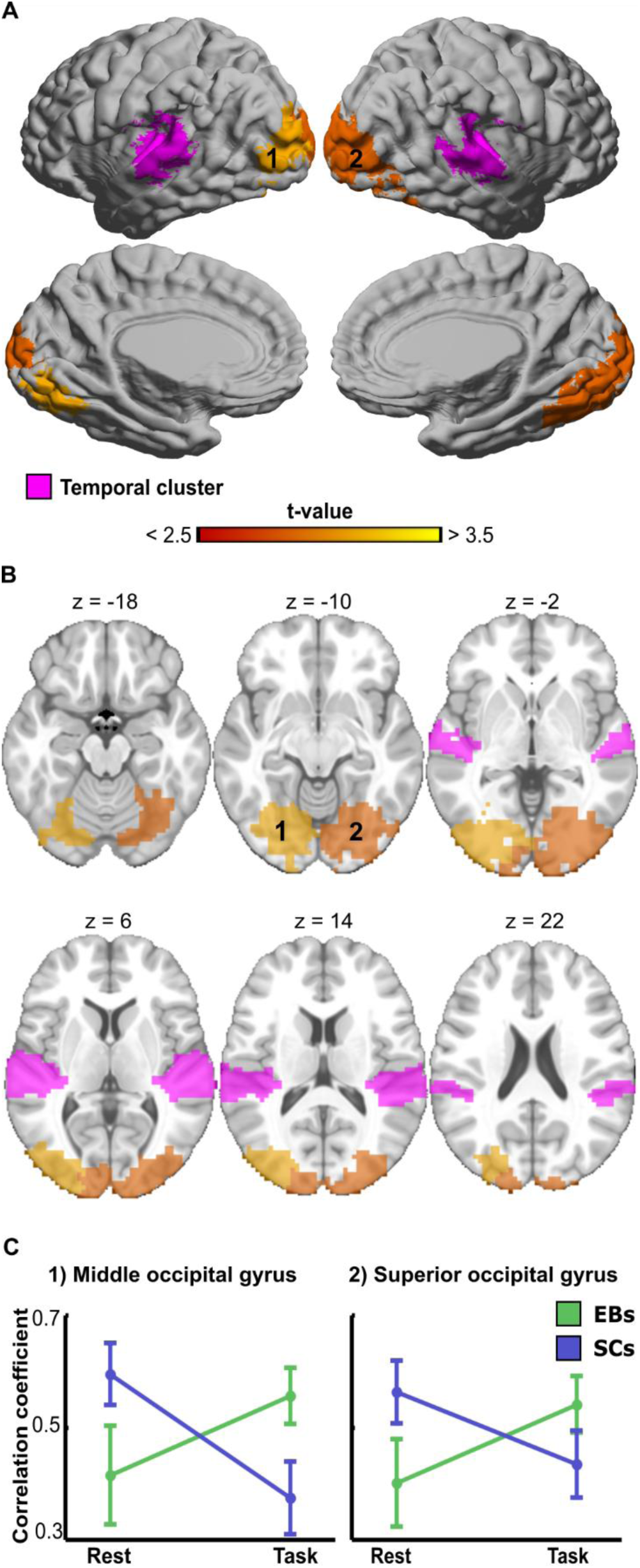
For the COI analysis, between-group differences in FC’s modulation effect [EBs(task – rest) vs SCs(task – rest)]. A, B) Maps show two regions, 1) middle occipital gyrus, and 2) superior occipital gyrus, for which the connectivity with the auditory regions was differently modulated by cognitive states in SCs versus EBs. Data is shown both on surfaces and brain slices. C) Plots show FC during rest and during the auditory task for both groups. Only significant interactions are plotted (q < 0.05, FDR corrected). Error bars represent standard errors.

### 3.2 Functional connectivity variability

Functional connections showing a distinct between-group modulation effect in the COI analysis also tended to be more variable across time in EBs, with the middle occipital gyrus being significantly more variable in EBs (p < 0.05) and the superior occipital gyrus showing a similar albeit non-significant trend (Fig. 5, see Supplementary Fig. 1 for all seven occipito-temporal connections). Moreover, a relationship between group differences in the modulation effect (EBs[task–rest] – SCs[task–rest]) and FC variability (EBs–SCs) is also supported by the highly significant correlation between these variables (*r*(6) = 0.9685, *p* < 0.001), with occipito-temporal connections showing stronger modulation effect in EBs also showing higher variability in EBs (Fig. 6). This high correlation cannot be taken as a mathematical artifact since it was specific to occipito-temporal connections but did not hold for the remainder of the brain’s FC (Fig. 6; *r*(1196) = −0.041, *p* = 0.16).

**Figure 5:**
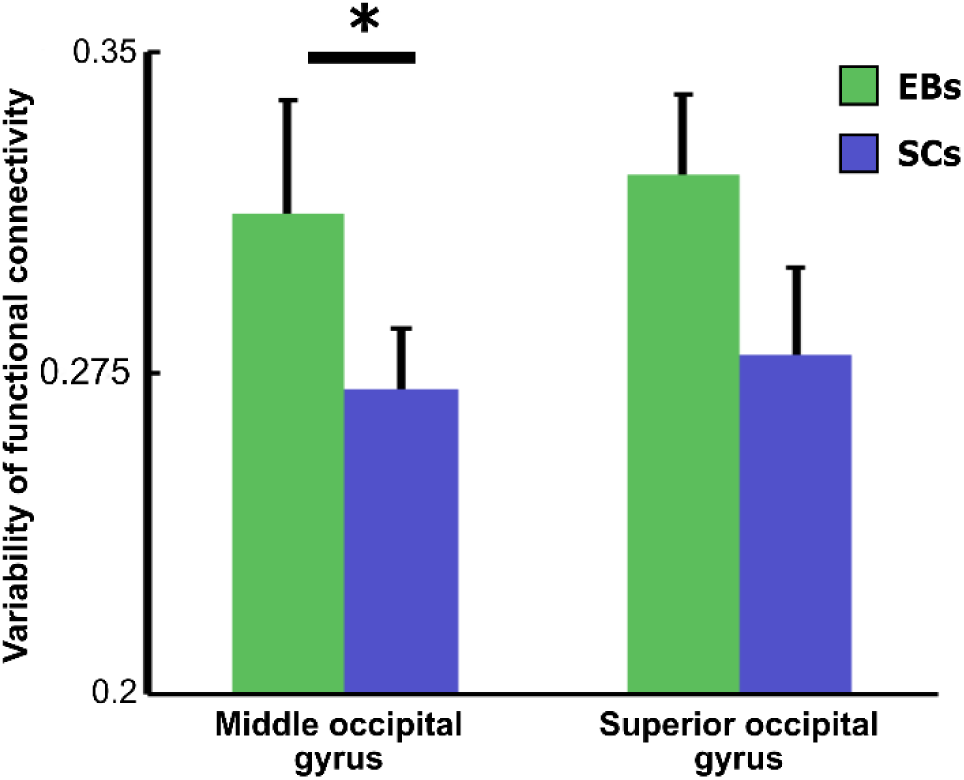
Results from the temporal variability (sliding windows) analysis of the resting-state run. Each column represents the variability in FC between the auditory region and one of two occipital regions. These are the same two connection which were differently modulated by cognitive states in EBs when compared to SCs. Error bars represent standard errors.* p < 0.05

**Figure 6:**
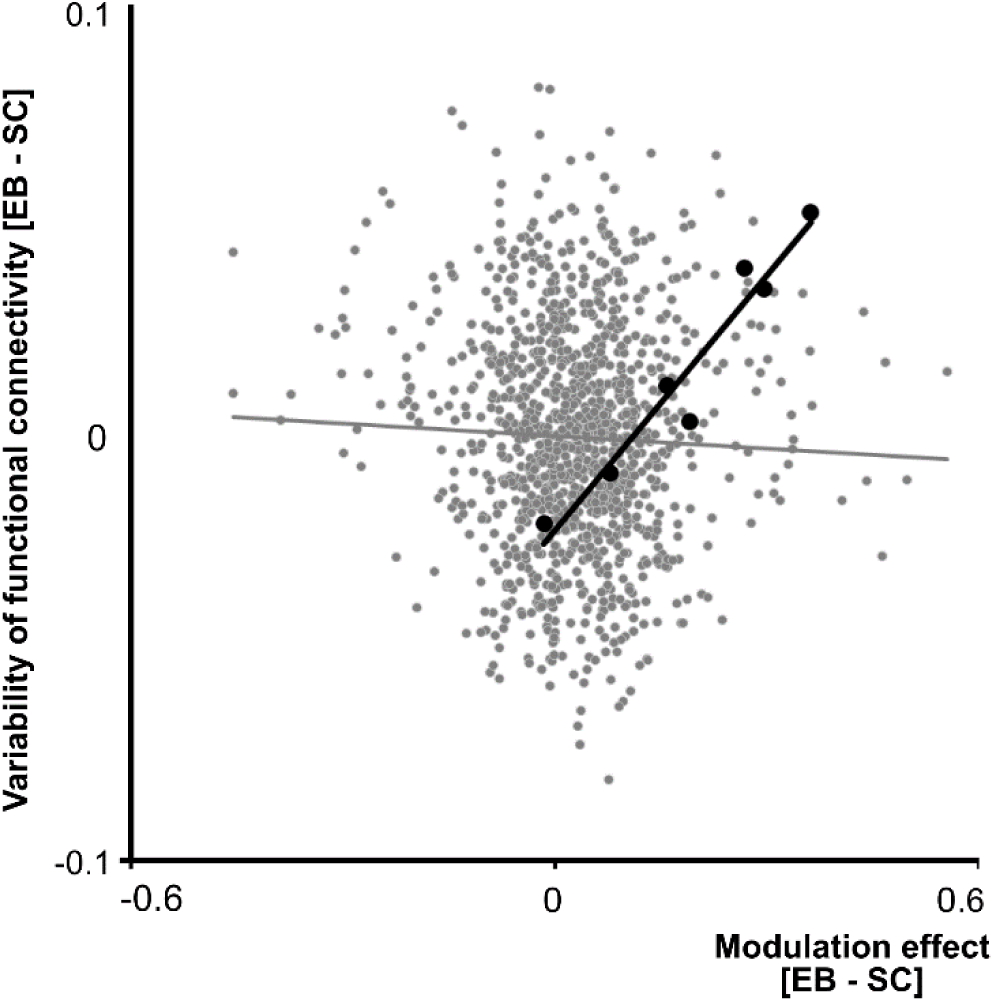
Correlation between the average difference in modulation across groups [EBs(Task – Rest) – SCs(Task – Rest)] and the average difference of FC variability across groups. Each black data point represents a single occipito-temporal functional connection (Correlation significant at *p* < 0.005). Each gray data point represent the strength of a single functional connection between two regions not found within the occipital mask (Correlation was non-significant, *p = 0.16*).

## 4 Discussion

In order to study how alterations of occipito-temporal connectivity due to early visual deprivation are influenced by cognitive state, EBs and SCs were scanned during rest and while involved in a demanding auditory task. Unbiased functional regions common across tasks and groups were defined (Bellec et al., 2010), and correlations between the region’s time series served as an index of FC which was compared across groups and cognitive states. We observed that occipito-temporal connectivity was lower during rest than during the auditory task in EBs, whereas the reverse pattern was observed in SCs (see Fig. 3). Our finding of an interaction effect between cognitive states and groups on FC is consistent with a body of recent studies showing how specific contexts of information processing fundamentally impact the strength of FC between brain regions (Eckert et al., 2008; Hasson et al., 2009; Moussa et al., 2011). For this reason, population differences in static RS networks metrics should not be considered to reliably index how networks instantiated during specific cognitive function differ across these populations.

Evidence of task-dependent connectivity modulation in SCs agrees with the observation of a decoupling between auditory and occipital regions during a visual task (Eckert et al., 2008). Crossmodal inhibition between auditory and visual cortices has been shown to take place during both auditory and visual stimulation (Laurienti et al., 2002). Clues as to the mechanisms underlying these results come from electrode recordings in mice revealing that auditory stimulation triggers inhibitory GABAergic release in V1 (Iurilli et al., 2012). In case of early blindness, it might be hypothesized that this inhibitory modulation of occipital regions triggered by sound switches to an excitatory state in the absence of competitive visual input during development (Collignon et al., 2009). Potential anatomical structures or physiological processes responsible for EB’s increased occipito-temporal connectivity include rewired or reinforced thalamo-cortical pathways as well as direct cortico-cortical connections between primary sensory cortices (Bavelier and Neville, 2002; Collignon et al., 2013; Karlen et al., 2006; Klinge et al., 2010; Shimony et al., 2006; Qin and Yu, 2013). It is worth noting here that sounds also has the potential to increase occipital activity even in sighted individuals in the context of audio-visual stimulation (Giard and Peronnet, 1999; Mercier et al., 2013; Rohe and Noppeney, In press) and that direct connection between primary auditory and visual cortices also exist in the absence of visual deprivation (Falchier et al. 2002, Rockland and Ojima, 2003). The observation of enhanced occipito-temporal connectivity in EBs during an auditory task is congruent with the plethora of task-dependent studies highlighting the functional involvement of EBs’ occipital cortex in auditory processing (Burton, 2003; Collignon et al., 2007; 2009; 2011; Weeks et al. 2000). Similarly, dynamic causal modeling studies show greater cortico-cortical connectivity between these regions in EBs than in SCs when participants are involved in auditory processing (Collignon et al., 2013; Fujii, 2009; Klinge et al., 2010). Finally, a recent magnetoencephalographic study showed that processing auditory inputs in blind produces stronger neural synchronization between auditory and visual cortices in the gamma band, suggesting that the deprived visual cortex is integrated into a larger network related to its new auditory function (Schepers et al., 2012).

At rest, FC between occipital and temporal regions tended to be lower for blind compared to sighted people, a finding that replicates several previous studies (see Bock and Fine, 2014 for review). What could explain EBs’ heightened occipito-temporal coupling during task while still shedding light on the reduced RSFC at rest? We hypothesized that one potential solution may reside in the fact that the occipital cortex of EBs involves in multiple independent functional modes. Evidence of cerebral regions involved in multiple overlapping functional modes exists, which notably manifests by the fact these regions switch membership between different temporally independent networks (Duncan and Owen 2000; Smith et al., 2012). This leads to temporally variable connectivity patterns for a specific region (see Hutchison et al., 2013a for a review), a perspective not addressed by using an entire fMRI run to compute of a single FC value per connection. By combining the contribution of all these modes, this single FC value obscures the true underlying functional organization of the brain (Hutchison et al., 2013b; Smith et al., 2012), likely underestimating the effective coupling between functional units. In EBs, the involvement of the occipital cortex in a large number of functional modes is already supported by studies showing its implications in various sensory and cognitive tasks including auditory and tactile processing as well as language, attention and episodic memory (Amedi et al., 2003, 2004; Bedny et al., 2011; Collignon et al., 2013; Burton, 2003; Gougoux et al., 2009; Noppeney, 2007; Pietrini et al., 2003; Raz et al., 2005). Therefore, lower occipito-temporal RSFC in EBs may result from the implication of their occipital lobe in a greater number of modes, attenuating FC while the brain is free to explore many modes (at rest), and unveiling the importance of these connections while modes are constrained (during a task). Based on the idea that, while at rest, cerebral regions involved in multiple modes would show increased FC variability across time with various networks (Allen et al., 2012), we measured the variability in occipito-temporal FC in both populations for the RS data. In agreement with our hypothesis, an occipito-temporal connection that was modulated by cognitive-states differently in EBs than in SCs also showed a higher variance in functional connectivity across time in EBs than SCs (Fig. 5). Interestingly, our data also supports the presence of this effect across all occipito-temporal connection’s since there was a strong and significant correlation between a connection’s temporal variability and its modulation effect (Fig. 6). Importantly, it supports the idea that partition in multiple modes (e.g. FC of high temporal variability), might shadow a region’s true potential within a network, which in turn can be unveiled during participation to specific tasks.

Another, but more speculative, possibility to explain lower occipito-temporal RSFC in EB, not mutually exclusive with the previous one, is that parts of EBs’ occipital cortex form a homogeneous network which is neither modality specific nor task specific. In SCs, studies have revealed the inferior frontal cortex, anterior cingulate cortex and inferior parietal sulci to present such characteristics and lead to the elaboration of the multiple-demand system theory (MD; Duncan and Owen, 2000, see Duncan 2010 for a review). According to this position, these regions’ lack of stimuli, modality, or task specificity makes them an optimal substrate for cognitive flexibility (Fedorenko et al., 2013). Coactivations of the MD system with a variety of task specific networks (Cabeza and Nyberg, 2000) are highly reminiscent of EBs occipital involvement in a collection of sensory and cognitive operations (see for review Frasnelli et al., 2011; Heimler et al., 2015; Ricciardi et al., 2014; Voss et al., 2011). This similarity between the MD system and EBs’ extends to how these regions respond to attentional demands. In SCs, fMRI studies show activations of MD regions to stimuli of various modalities when they are attended but not when they are unattended (Hon et al., 2006). Similarly, occipital responses to auditory and tactile stimuli have been shown to be dependent upon the attentional focus in EBs (Kujala et al., 1995; 2005; Sadato et al., 1996; Weaver and Stevens, 2007). However, this is also true of early sensory areas (Kastner et al., 1999). Finally, further support as to the involvement of EBs occipital lobe with the MD system comes from a number of RSFC studies which evidence stronger coupling between frontal areas, which seem congruent with those belonging to the MD system and executive functions, and the visual cortices (Bedny et al., 2010; 2011; Burton et al., 2014; Deen, et al., 2015; Liu et al., 2007; Wang et al., 2013). Still, the true hallmark of the MD system is that its neurons rapidly reorient their response tuning to task relevant features (Duncan, 2010), a fact which has, to our knowledge, never been investigated in EBs. Thus, though the similarities between the MD system and EBs’ occipital lobes may be seductive, further research is required to investigate this point.

Our results therefore open new avenues of research on the connectivity of the occipital cortex of the blind and raise attention on the fact that group differences in connectivity measures inferred from resting-state do not readily generalize to specific task-dependent brain-state (e.g. auditory processing). In particular, our study provides a simple foundation for analyses of state-dependent changes in connectivity in visually deprived individuals. The importance of our results goes beyond the study of blindness by extending a flourishing number of studies (Çetin et al., 2014; Nair et al., 2014) highlighting how inferring the functional connectivity profile of a region solely based on RS data is subject to caution.

## Supplementary tables

**Supplementary table 1.**
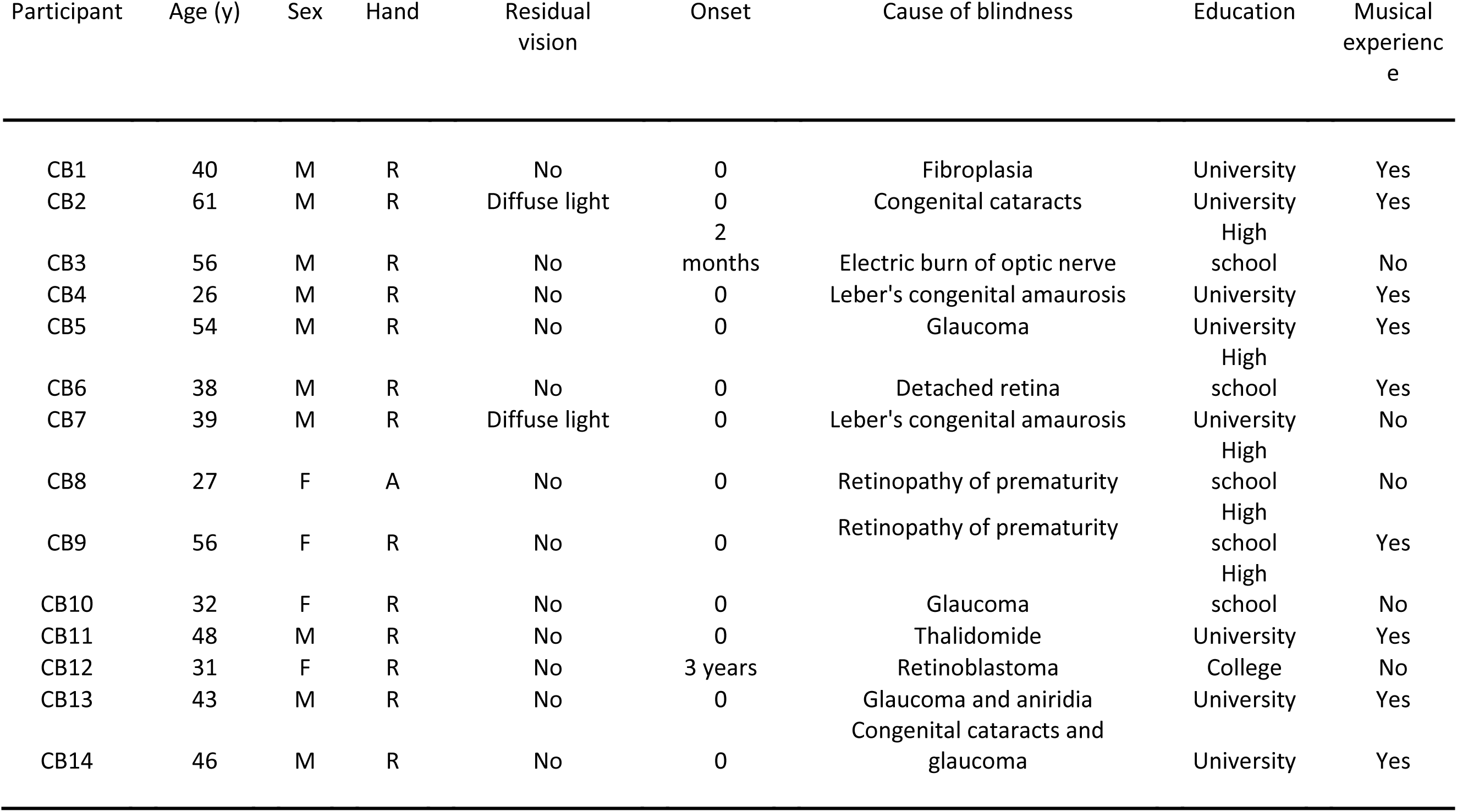
Characteristics of the blind participants.

**Supplementary table 2.** Central coordinates and average radius in standard deviation of temporal and occipital networks at resolution 50.

| Network number<br>and region's name | Left hemisphere |  |  |  | Right hemisphere |  |  |  |
| --- | --- | --- | --- | --- | --- | --- | --- | --- |
|  | x | y | z | Approximate<br>radius(mm,sd) | x | y | z | Approximate<br>radius(mm,sd) |
| 25. Superior temporal<br>gyrus | -52 | -23 | 9 | 5,00 | 55 | -25 | 11 | 4,94 |
| 37. Middle occipital gyrus | -28 | -81 | 2 | 5,80 | - | - | - | - |
| 38.Cuneus | -12 | -81 | 30 | 4,85 | 16 | -82 | 28 | 4,56 |
| 42. Calcarine fissure | -13 | -67 | 6 | 4,34 | 14 | -68 | 6 | 4,78 |
| 43. Lingual gyrus | -25 | -86 | -16 | 7,55 | 30 | -80 | -20 | 9,20 |
| 46. Middle temporal gyrus | -42 | -68 | 19 | 8,81 | 47 | -64 | 16 | 7,53 |
| 47. Superior occipital<br>gyrus | -11 | -95 | 10 | 3,73 | 26 | -82 | -1 | 7,32 |
| 50. Inferior temporal gyrus | -50 | -42 | -17 | 7,28 | 56 | -37 | -18 | 9,49 |

**Supplementary table 3.** Average and standard deviation of fisher transform of correlation coefficient for each occipito-temporal functional connection investigated.

| Functional connectivity<br>between temporal cluster<br>(#25) and: | Early blind participants |  |  |  |  |  | Sighted controls |  |  |  |  |  |
| --- | --- | --- | --- | --- | --- | --- | --- | --- | --- | --- | --- | --- |
|  | Rest |  | Task<sup>1</sup> |  | Task<sup>2</sup> |  | Rest |  | Task<sup>1</sup> |  | Task<sup>2</sup> |  |
|  | M | SD | M | SD | M | SD | M | SD | M | SD | M | sd |
| 37. Middle occipital gyrus | 0,416 | 0,080 | 0,557 | 0,052 | 0,564 | 0,056 | 0,595 | 0,048 | 0,375 | 0,059 | 0,400 | 0,065 |
| 38. Cuneus | 0,594 | 0,053 | 0,700 | 0,071 | 0,702 | 0,077 | 0,585 | 0,048 | 0,393 | 0,070 | 0,413 | 0,076 |
| 42. Calcarine fissure | 0,535 | 0,075 | 0,725 | 0,052 | 0,727 | 0,061 | 0,546 | 0,069 | 0,658 | 0,054 | 0,673 | 0,057 |
| 43. Lingual gyrus | 0,211 | 0,065 | 0,370 | 0,045 | 0,377 | 0,047 | 0,395 | 0,061 | 0,396 | 0,032 | 0,405 | 0,029 |
| 46. Middle temporal gyrus | 0,638 | 0,082 | 0,635 | 0,081 | 0,632 | 0,081 | 0,704 | 0,055 | 0,510 | 0,067 | 0,532 | 0,073 |
| 47. Superior occipital gyrus | 0,402 | 0,079 | 0,542 | 0,053 | 0,543 | 0,056 | 0,564 | 0,03 | 0,435 | 0,054 | 0,456 | 0,060 |
| 50. Inferior temporal gyrus | 0,444 | 0,094 | 0,382 | 0,084 | 0,395 | 0,084 | 0,450 | 0,073 | 0,404 | 0,049 | 0,424 | 0,052 |

<sup>1</sup> Task run with the effect of block modeled out of the time series.
<sup>2</sup> Task run with basic preprocessing.

## Supplementary figure captions

**Supplementary figure 1.**
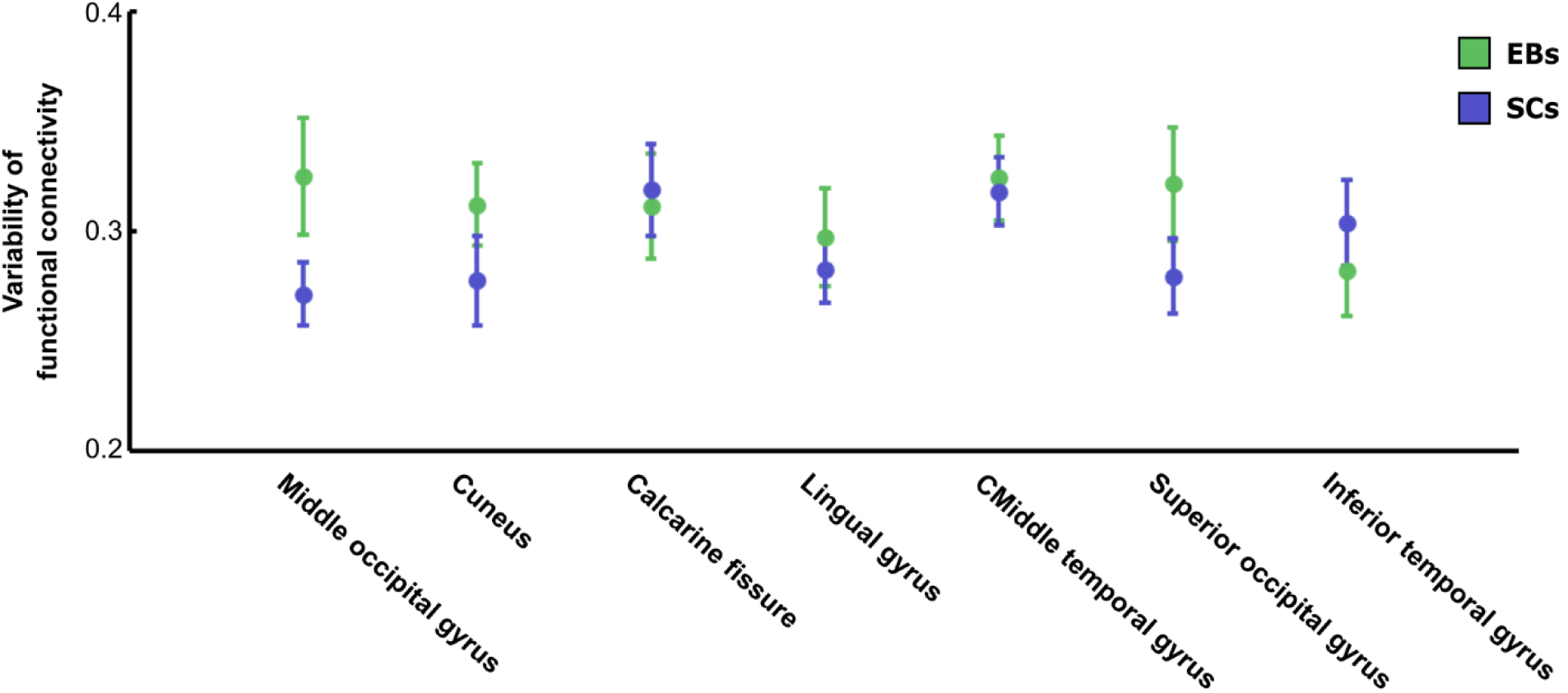
Group average and standard error of FC temporal variability at rest for each functional connection between the temporal region and the 7 occipital regions.

## References

Allen EA. Damaraju E. Plis SM. Erhardt EB. Eichele T. Calhoun. VD. 2012. Tracking Whole–Brain Connectivity Dynamics in the Resting State. Cereb Cortex. 24:663–676.

Amedi A. Floel A. Knecht S. Zohary E. Cohen LG. 2004. Transcranial magnetic stimulation of the occipital pole interferes with verbal processing in blind subjects. Nat Neurosci. 7:1266–1270.

Amedi A. Raz N. Pianka P. Malach R. Zohary E. 2003. Early 'visual' cortex activation correlates with superior verbal memory performance in the blind. Nat Neurosci. 6:758–766.

Andric M. Hasson U. 2015. Global features of functional brain networks change with contextual disorder. Neuroimage. 117:103–113.

Bartels A. Zeki S. 2005. Brain dynamics during natural viewing conditions–A new guide for mapping connectivity in vivo. Neuroimage. 24:339–349.

Bavelier D. Neville HJ. 2002. Cross-modal plasticity: where and how? Nat Rev Neurosci. 3:443–452.

Beckmann CF. DeLuca M. Devlin JT. Smith SM. 2005. Investigations into resting–state connectivity using independent component analysis. Philos Trans R Soc Lond B Biol Sci. 360:1001–1013.

Bedny M. Konkle T. Pelphrey K. Saxe R. Pascual–Leone A. 2010. Sensitive Period for a Multimodal Response in Human Visual Motion Area MT/MST. Curr Biol. 20:1900–1906.

Bedny M. Pascual–Leone A. Dodell–Feder D. Fedorenko E. Saxe R. 2011. Language processing in the occipital cortex of congenitally blind adults. Proc Natl Acad Sci U S A. 108:4429–4434.

Bellec P. Benhajali Y. Carbonell F. Dansereau C. Albouy G. Pelland M. Orban P. 2015. Impact of the resolution of brain parcels on connectome–wide association studies in fMRI. Neuroimage. 123:212–228.

Bellec P. Perlbard V. Jbabdi S. Pélégrini–Issac M. Anton JL. Doyon J. Benali H. 2006. Identification of large–scale networks in the brain using fMRI. Neuroimage. 29:1231–1243.

Bellec P. Rosa–Neto P. Lyttelton O. Benali H. Evans AC. 2010. Multi–level bootstrap analysis of stable clusters in resting–state fMRI. Neuroimage. 51:1126–1139.

Bellec P. Lavoie–Courchesne L. Dickinson P. Lerch JP. Zijdenbos AP. Evans AC. 2012. The pipeline system for Octave and Matlab PSOM: a lightweight scripting framework and execution engine for sceintific workflows. Front Neuroinform. 6: 1–18.

Benjamini Y. Hochberg Y. 1995. Controlling the false–discovery rate: a practical and powerful approach to multiple testing. J R Stat Soc Series B Stat Methodol. 57:289–300.

Biswal B. Yetkin FZ. Haughton VM. Hyde JS. 1995. Functional connectivity in the motor cortex of resting human brain using echo–planr MRI. Magn Reson Med. 34:537–541.

Bock AS. Fine I. 2014. Anatomical and Functional Plasticity in Early Blind Individuals and the Mixture of Experts Architecture. Front Hum Neurosci. 8:971.

Burton H. 2003. Visual cortex activity in early and late blind people. J Neurosci. 23:4005–4011.

Burton H. Snyder A. Raichle M. 2014. Resting state functional connectivity in early blind humans. Front Syst Neurosci. 8:51.

Cabeza R. Nyberg L. 2000. Imaging cognition II: An empirical review of 275 PET and fMRI studies. J Cogn Neurosci. 12:1–47.

Çetin MS. Christensen F. Abbott CC. Stephen JM. Cañive JM. Bustillo JR. Pearlson GD. Clahoun. V.D. 2014. Thalamus and posterior temporal lobe show greater inter–network connectivity at rest and across sensory paradigms in schizophrenia. Neuroimage. 97:117–126.

Cohen LG. Celnik P. Pascual-Leone A. Corwell B. Falz L. Dambrosia J. Honda M. Gerloff C. Catalá MD. et al. 1997. Functional relevance of cross–modal plasticity in blind humans. Nature. 389:180–183.

Collignon O. Davare M. Olivier E. De Volder AG. 2009. Reorganisation of the right occipito–parietal stream for auditory spatial processing in early blind humans. A transcranial magnetic stimulation study. Brain Topogr. 21:232–240.

Collignon O. Dormal G. Albouy G. Vandewalle G. Voss P. Phillips C. Lepore F. 2013. Impact of blindness onset on the functional organization and the connectivity of the occipital cortex. Brain. 136:2769–2783.

Collignon O. Lassonde M. Lepore F. Bastien D. Veraart C. 2007. Functional cerebral reorganization for auditory spatial processing and auditory substitution of vision in early blind subjects. Cereb Cortex. 17: 457–465.

Collignon O. Vandewall G. Voss P. Albouy G. Charbonneau G. Lassonde M. Lepore F. 2011. Functional specialization for auditory–spatial processing in the occipital cortex of congenitally blind humans. Proc Natl Acad Sci U S A. 108:4435–4440.

Cordes D. Haughton VM. Arfanakis K. Wendt GJ. Turski PA. Moritz CH. Quigley MA. Meyerand. M.E. 2000. Mapping functionally related regions of brain with functional connectivity MR imaging. AJNR Am J Neuroradiol. 21:1636–1644.

Debas K. Carrier J. Barakat M. Marrelec G. Bellec P. Tahar AH. Karni A. Ungerleider LG.Benali H. Doyon J. 2014. Off-line consolidation of motor sequence learning results in greater integration within a cortico-striatal functional network. NeuroImage. 99:50–58.

Deen B, Saxe R. Bedny M. (2015). Occipital Cortex of Blind Individuals Is Functionally Coupled with Executive Control Areas of Frontal Cortex. Journal of Cognitive Neurosciences 27:1633–1647.

Deco G. Jirsa VK. & McIntosh AR. (2011). Emergin concepts for the dunamical organization of resting-state activity in the brain. Nature reviews: neuroscience, 12, 43–56.

Di X. Gohel S. Kim EH. Biswal BB. 2013. Task vs. rest – different network configurations between the coactivation and the resting-state brain networks. Front Hum Neurosci. 7:493.

Dormal G. Collignon O. 2011. Functional selectivity in sensory–deprived cortices. J Neurophysiol. 105:2627–2630.

Duncan J. 2010. The multiple–demand MD system of the primate brain: mental pograms for intelligent behaviour. Trends Cogn Sci. 14:172–179.

Duncan J. Owen AM. 2000. Common regions of the human frontal lobe recruited by diverse cognitive demands. Trends Neurosci. 23:475–483.

Eckert MA. Kamdar NV. Chang CE. Beckmann CF. Greicius MD. Menon V. 2008. A Cross–Modal System Linking Primary Auditory and Visual Cortices: Evidence From Intrinsic fMRI Connectivity Analsysis. Hum Brain Mapp. 29:848–857.

Fair DA. Schlaggar BL. Cohen AL. Miezin FM. Dosenbach NU. Wenger KK. Fox MD. Snyder AZ Raichle ME. Petersen SE. 2007. A method for using blocked and evern–related fMRI data to study “resting state” functional connectivity. Neuroimage. 35:396–405.

Falchier A. Clavagnier S. Barone P. Kennedy H. 2002. Anatomical evidence of multimodal integration in primate striate cortex. J Neurosci. 22:5749–5759.

Fedorenko E. Duncan J. Kanisher N. 2013. Broad domain generality in focal regions of frontal and parietal cortex. Proc Natl Acad Sci U S A. 110:16616–16621.

Fonov VS. Evans AC. Botteron K. Almli CR. McKinstry RC. Collins DL BDCG. 2011. Unbiased average age-appropriate atlases for pediatric studies. Neuroimage. 54:313–317.

Fonov VS. Evans AC. McKinstry RC. Almli CR. Collin DL. 2009. Unbiased nonlinear average age appropriate brain template from birth to adulthood. Neuroimage. 47:s102.

Frasnelli J. Collignon O. Voss P. Lepore F. 2011. Sensory Rehabilitation in the Plastic Brain. Prog Brain Res. 191:211–231.

Friston K. 1994. Functional and Effective Connectivity in Neuroimaging: A Synthesis. Hum Brain Mapp. 2:56–78.

Friston KJ. Frith CD. Liddle PF. Frackowiak RS. 1993. Functional connectivity: the principal–component analysis of large PET data sets. J Cereb Blood Flow Metab. 13:5–14.

Friston K. Harrison L. Penny W. 2003. Dynamic causal modelling. Neuroimage. 19:1273–1302.

Fuji T. Tanabe HC. Kochiyama T. Sadato N. 2009. An investigation of cross–modal plasticity of effective connectivity in the blind by dynamic causal modeling of functional MRI data. Neurosci Res. 65:175–186.

Giard MH. Peronnet F. 1999. Auditory-visual integration during multimodal object recognition in humans: a behavioral and electrophysiological study. J Cogn Neurosci. 11:473–490.

Gordon EM. Breeden AL. Bean SE. Vaidya CJ. 2014. Working memory–related changes in functional connectivity persist beyond task disengagement. Hum Brain Mapp. 35:1004–1017.

Gougoux F. Belin P. Voss P. Lepore F. Lassonde M. Zatorre RJ. 2009. Voice perception in blind persons: a functional magnetic resonance imaging study. Neuropsychologia. 47:2967–2974.

Gougoux F. Zatorre RJ. Lassonde M. Voss P. Lepore F. 2005. A functional neuroimaging study of sound localization: visual cortex activity predicts performance in early–blind individuals. PLoS Biol. 3:27.

Hampson M. Peterson BS. Skudlarski P. Gatenby JC. Gore JC. 2002. Detection of Functional Connectivity Using Temporal Correlations in MR Images. Hum Brain Mapp. 15:247–262.

Hampson M. Olson IR. Leung H–C. Skudlarski P. Gore JC. 2004. Changes in functional connectivity of human MT/V5 with visual motion input. NeuroReport. 15:1315–1319.

Heimler B. Striem–Amit E. Amedi A. 2015. Origins of task–specific sensory–independent organization in the visual and auditory brain: neuroscience evidence. open questions and clinical implications. Curr Opin Neurobiol. 35:169–177.

Hölig C. Föcker J. Best A. Röder B. Büchel C. 2014. Crossmodal plasticity in the fusiform gyrus of late blind individuals during voice recognition. Neuroimage. 103:374–382.

Hasson U. Nusbaum HC. Small SL. 2009. Task–dependent organization of brain regions active during rest. Proc Natl Acad Sci U S A. 106:10841–10846.

Hindriks R. Adhikari MH. Murayama Y. Ganzetti M. Mantini D. Logothetis NK. Deco G. 2016. Can sliding-window correlations reveal dynamic functional connectivity in resting-state fMRI? Neuroimage. 127:242–356.

Hon N. Epstein RA. Owen AM. Duncan J. 2006. Frontoparietal activity with minimal decision and control. J Neurosci. 26:9805–9809.

Hutchison RM. Womelsdorf T. Allen E. Bandettini PA. Calhoun V. Corbetta M. Della Penna S. Duyn JH. Glover GH. Gonzalez-Castillo J. et al. 2013a. Dynamic functional connectivity: Promises. issues. and interpretations. Neuroimage. 80:360–378.

Hutchison RM. Womelsdorf T. Gati JS. Everling S. Menon RS. 2013b. Resting–state networks show dynamic functional connectivity in awake humans and anesthetized macaques. Hum Brain Mapp. 34:2154–2177.

Iurilli G. Ghezzi D. Olcese U. Lassi G. Nazzaro C. Tonin R. Tucci V. Benfenati F. Medini P. 2012. Sound–Driven Synaptic Inhibition in Visual Cortex. Neuron. 73:814–828.

Jiang T. He Y. Zang Y. Weng X. 2004. Modulation of Functional Connectivity During the Resting State and the Motor Task. Hum Brain Mapp. 22:63–71.

Karahanoğlu FI. Caballero-Gaudes C. Lazeyras F. Van de Ville D. (2013). Total activation: fMRI deconvolution through spatio-temporal regularization. Neuroimage, 73, 121–134.

Karlen SJ. Kahn DM. Krubitzer L. 2006. Early blindness results in abnormal corticocortical and thalamocortical connections. Neuroscience. 142:843–858.

Kastner S. Pinsk MA. De Weerd P. Desimone R. Ungerleider LG. 1999. Increased activity in human visual cortex during directed attention in the absence of visual stimulation. Neuron, 22:751–761.

Klinge C. Eippert F. Röder B. Büchel C. 2010. Corticocortical Connections Mediate Primary Visual Cortex Responses to Auditory Stimulation in the Blind. J Neurosci. 30:12798–12805.

Kujala T. Huotilainen M. Sinkkonen J. Ahonen AI. Alho K. Hämäläinen MS. Ilmoniemi FJ. Kajola M. Knuutila JE. Lavikainen J. 1995. Visual cortex activation in blind humans during sound discrimination. Neurosci Lett. 183:143–146.

Kujala T. Plava MJ. Salonen O. Alku P. Huotilainen M. Järvinen A. Näätänen R. 2005. The role of blind humans’ visual cortex in auditory change detection. Neurosci Lett. 379:127–131.

Laird A. Eickhoff SB. Rottschy C. Bzdok D. Ray KL. Fox PT. 2013. Networks of Task Co-Activations. Neuroimage. 80:505–514.

Laurienti. PJ. Burdette JH. Wallace MT. Yen Y–F. Field AS. Stein BE. 2002. Deactivation of Sensory–Specific Cortex by Cross–Modal Stimuli. J Cogn Neurosci. 14:420–429.

Leclerc C. Saint–Amour D. Lavoie ME. Lassonde M. Lepore F. 2000. Brain functional reorganization in early blind humans revealed by auditory event–related potentials. Neuroreport. 11:545–550.

Lewis CM. Baldassarre A. Committeri G. Romani GL. Corbetta M. 2009. Learning sculpts the spontaneous activity of the resting human brain. Proc Natl Acad Sci U S A. 106:17558–17563.

Logothetis, N.K., & Wandell, B.A. (2004). Interpreting the BOLD Signal. Annual Review of Physiology, 66, 735–769.

Liu C. Liu Y. Li W. Wang D. Jiang T. Zhang Y. Yu C. 2011. Increased regional homogeneity of blood oxygen level–dependent signals in occipital cortex of early blind individuals. Neuroreport. 22:190–194.

Liu Y. Yu C. Liang M. Li J. Tian L. Zhou Y. Qin W. Li K. Jiang T. 2007. Whole brain functional connectivity in the early blind. Brain. 130:2085–2096.

Mennes M. Kelly C. Colcombe S. Castellanos FX. Milham MP. 2013. The extrinsic and intrinsic functional archtectures of the human brain are not equivalent. Cereb Cortex. 23:223–229.

Mercier MR. Foxe JJ. Fiebelkorn IC. Butler JS. Schwartz TH. Molholm S. 2013. Auditory-driven phase reset in visual cortex: human electrocorticography reveals mechanisms of early multisensory integration. Neuroimage. 79:19–29.

Moussa MN. Vechlekar CD. Burdette JH. Steen MR. Hugenschmidts CE. Laurienti PJ. 2011. Changes in cognitive state alter human functional brain networks. Front Hum Neurosci. 5:83.

Nair A. Keown CL. Datko M. Shih P. Keehn B. Müller R–A. 2014. Impact of Methodological Variables on Functional Connectivity Findings in Autism Spectrum Disorders. Hum Brain Mapp. 35:4035–4048.

Newton AT. Morgan VL. Gore JC. 2007. Task Demand Modulation of Steady–State Functional Connectivity to Primary Motor Cortex. Hum Brain Mapp. 28:663–672.

Nir Y. Hasson U. Levy I. Yeshurun Y. Malach R. 2006. Widespread functional connectivity and fMRI fluctuations in human visual cortex in the absence of visual stimuation. Neuroimage. 30:1313–1324.

Noppeney U. 2007. The effects of visual deprivation on functional and structural organization of the human brain. Neurosci Biobehav Rev. 31:1169–1180.

Orban P. Doyon J. Petrides M. Mennes M. Hoge R. Bellec P. 2015. The Richness of Task-Evoked Hemodynamic Responses Defines a Pseudohierarchy of Functionally Meaningful Brain Networks. Cereb Cortex. 25:2658–2669.

Park B. Ko JH. Lee JD. Park HJ. 2013. Evaluation of node–inhomogeneity effects on the functional brain network properties using an anatomy–constrained hierarchical brain parcellation. PLoS One. 8:e74935. doi:10.1371/journal.pone.0074935. eCollection 2013.

Pietrini P. Furey ML. Ricciardi E. Gobbini MI. Wu WH. Cohen L. Guazzelli M. Haxby J.V. 2004. Beyond sensory images: Object–based representation in the human ventral pathway. Proc Natl Acad Sci U S A. 101:5658–5663.

Power JD. Barnes KA. Snyder AZ. Schlaggar BL. Petersen S.E. 2012. Spurious but systematiccorrelations in functional connectivity MRI networks arise from subject motion. Neuroimage. 59:2142–2154.

Ptito M. Fumal A. de Noordhout AM. Schoenen J. Gjedde A. Kupers R. 2008. TMS of the occipital cortex induces tactile sensations in the fingers of blind Braille readers. Exp Brain Res. 184:193–200.

Qin W. Yu C. 2013. Neural Pathways Conveying Novisual Information to the Visual Cortex. Neural Plast. 2013.864920:doi:10.1155/2013/864920

Raz N. Amedi A. Zohary E. 2005. V1 activation in congenitally blind humans is associated with episodic retrieval. Cereb Cortex. 15:1459–1468.

Reich L. Maidenbaum S. Amedi A. 2012. The brain as a flexible task machine: implications for visual rehabilitation using noninvasive vs. invasive approaches. Curr Opin Neurol. 25:86–95.

Ricciardi E. Bonino D. Pellegrini S. Pietrini P. 2014. Mind the blind brain to understand the sighted one! Is there a supramodal cortical functional architecture? Neurosci Biobehav Rev. 41:64–77.

Rockland KS. Ojima H. 2003. Multisensory convergence in calcarine visual areas in macaque monkey. Int J Psychophysiol. 50:19–26.

Röder B. Teder–Sälejärvi W. Sterr A. Rösler F. Hillyard SA. Neville H. 1999. Improved auditory spatial tuning in blind humans. Nature. 400:162–166.

Rohe T. Noppeney U. In press. Distinct Computational Principles Govern Multisensory Integration in Primary Sensory and Association Cortices. Curr Biol. doi:10.1016/j.cub.2015.12.056

Ryali S. Chen T. Padmanabhan A. Cai W. Menon V. 2015. Development and validation of consensus clustering-based framework for brain segmentation using resting fMRI. J Neurosci Methods. 240:128–140.

Sadato N. Pascual-Leone A. Grafman J. Ibañez V. Deiber MP. Dold G. Hallett M. 1996. Nature. 380:526–528.

Schepers IM. Hipp JF. Schneider TR. Röder B. Engel AK. 2012. Functionally specific oscillatory activity correlates between visual and auditory cortex in the blind. Brain. 135:922–934.

Shimony JS. Burton H. Epstein AA. McLaren DG. Sun SW. Snyder AZ. 2006. Diffusion tensor imaging reveals white matter reorganization in early blind humans. Cereb Cortex. 16:1653–1661.

Shirer WR. Ryali S. Rykhlevskaia E. Menon V. Greicius MD. 2012. Decoding subject–driven cognitive states with whole–brain connectivity patterns. Cereb Cortex. 22:158–165.

Smith SM. Fox PT. Miller KL. Glahn DC. Fox PM. Mackay CE. Filippini N. Watkins KE. Toro R. Laird AR. Beckmann CF. 2009. Correspondence of the brain’s functional architecture during activation and rest. Proc Natl Acad Sci U S A. 106:13040–13045.

Smith SM. Miller KL. Moeller S. Xu J. Auerbarch EJ. Woolrich MW. Beckmann CF. Jenkinson M. Andersson J. Glasser MF et al. 2012. Proc Natl Acad Sci U S A. 109:3131–3136.

Souza, P., Gehani, N., Wright, R., McCloy, D. (2013). The advantage of knowing the talker. Journal of the American Academy of Audiology, 24(8), 689–700.

Striem-Amit E. Ovadia-Caro S. Caramazza A. Margulies S. Villringer A. Amedi A. 2015. Functional connectivity of visual cortex in the blind follows retinotopic organization principles. Brain. 138:1679–1695.

Tambini A. Ketz N. Davachi L. 2010. Enhanced brain correlations during rest are related to memory for recent experiences. Neuron. 65:280–290.

van den Heuvel MP. Hulshoff Pol HE. 2010. Exploring the brain network: a review on resting–state fMRI functional connectivity. Eur Neuropsychopharmacol. 20:519–534.

Voss P. Collignon O. Lassonde M. Lepore F. 2011. Adaptation to sensory losss. Wiley Interdiscip Rev Cogn Sci. 2:238. doi:10.1002/wcs.139.

Wang D. Qin W. Liu Y. Zhang Y. Jiang T. Yu C. 2014. Altered resting–state network connectivity in congenital blind. Hum Brain Mapp. 35:2573–2581.

Weaver KE. Steven AA. 2007. Attention and Sensory Interactions within the Occipital Cortex in the Early Blind: An fMRI Study. J Cogn Neurosci. 19:315–330.

Weeks R. Horwitz B. Aziz-Sultan A. Tian B. Wessinger CM. Cohen LG. Hallett M. Rauschecker JP. 2000. A positron emission tomographic study of auditory localization in the congenitally blind. J Neurosci. 20:2664–2672.

Wig GS. Laumann TO. Petersen SE. 2014. An approach for parcellating human cortical areas using resting–sate correlations. Neuroimage. 93:276–291.

Wittenberg GF. Werhahn KJ. Wassermann EM. Herscovitch P. Cohen LG. 2004. Functional connectivity between somatosensory and visual cortex in early blind humans. Eur J Neurosci.. 20:1923–1927.

Yu C. Liu Y. Li J. Zhou Y. Wang K. Tian L. Qin Q. Jiang T. Li. K. 2008. Altered Functional Connectivity of Primary Visual Cortex in Early Blindness. Hum Brain Mapp. 29:533–543.

Zalesky A. Fornito A. Harding IH. Cocchi L. Yücel M. Pantelis C. Bullmore ET. 2010. Whole-brain anatomical networks: does the choice of nodes matter? Neuroimage. 50:970–983.

